# Computational Translation of Mouse Models of Osteoarthritis Predicts Human Disease

**DOI:** 10.1101/2025.02.23.639777

**Authors:** Maya R. Frost, Brendan K. Ball, Meghana Pendyala, Stephen R. Douglas, Douglas K. Brubaker, Deva D. Chan

**Affiliations:** Weldon School of Biomedical Engineering, Purdue University; Department of Biomedical Engineering, Rensselaer Polytechnic Institute; School of Mechanical Engineering, Purdue University; Center for Global Health and Diseases, Department of Pathology, School of Medicine, Case Western Reserve University; Blood Heart Lung Immunology Research Center, University Hospitals Cleveland Medical Center

**Keywords:** Osteoarthritis, translational modeling, transcriptomics Running title: Osteoarthritis cross-species translation

## Abstract

**Objective:** Translation of biological insights from preclinical studies to human disease is a pressing challenge in biomedical research, including in osteoarthritis. Translatable Components Regression (TransComp-R) is a computational framework that has previously been used to synthesize preclinical and human OA data to identify biological pathways predictive of human disease conditions. We aimed to evaluate the translatability of two common murine models of post-traumatic osteoarthritis – surgical destabilization of the medial meniscus (DMM) and noninvasive anterior cruciate ligament rupture (ACLR) – to transcriptomics cartilage data from human OA outcomes.

**Design:** Transcriptomics cartilage data of DMM and ACLR mouse and human data was acquired from Gene Expression Omnibus. TransComp-R was used to project human OA data into a mouse model (DMM or ACLR) principal component analysis space. The principal components (PCs) were regressed against human OA conditions using increasing complexity of linear regression models incorporating human demographic covariates of OA, sex, and age. Biological pathways of the mouse PCs that significantly stratified human OA and control groups were then interpreted using Gene Set Enrichment Analysis.

**Results:** From the TransComp-R model, we identified different enriched biological pathways across DMM and ACLR models. While PCs among the DMM models revealed pathways associated with cell signaling and metabolism, ACLR PCs represented immune function and cellular pathways associated with OA condition. The immune pathways presented in the ACLR further highlighted the potential relevance of the OA pathways observed in human conditions.

**Conclusions:** The ACLR mouse model more successfully predicted human OA conditions, particularly with the human control groups without a history of joint injury or disease. Cross-species translational approaches support the selection of preclinical models intended for therapeutic discovery and pathway analysis in humans.

## INTRODUCTION

Osteoarthritis (OA) affects an estimated 250 million people worldwide, and its prevalence continues to rise as obesity rates increase and the population ages.^1^ Animal models of OA are necessary to study the mechanisms underlying the development and progression of OA and to evaluate the efficiency of disease interventions.^2^ Although murine models enable careful manipulation of experimental conditions and biological variables, mice differ in size, anatomy, and biomechanics from humans and demonstrate differences in disease progression,^2–6^ timeline of disease,^7, 8^ demographic variability,^6, 9, 10^ and disease heterogeneity. ^11^ While these descriptive differences have been cataloged, comparative approaches can be misleading for translatability.^12–14^ Therefore, predictive methods to overcome these barriers are needed to relate mouse to human OA and develop translatable, efficacious interventions.

Indeed, the translation of biological insights from animal models to humans remains a foundational challenge in preclinical biomedical research.^15, 16^ Clinical trial failure of inflammatory modulatory drugs is potentially attributable in part to differences in mouse and human immunology.^12–14^ A growing body of comparative cross-species studies^17^ that leverage study of the homology of mouse and human genomes, epigenomes, and transcriptomes^18^ has potential to improve the use and interpretation of animal models. Such efforts could guide future use of animal models to better understand characteristics and treatment of human OA.^19^ However, no studies to date have systematically interrogated these cross-species relationships in the context of biological signature of OA mouse models mapped to those of OA human patients. We and others have developed computational systems biology-based approaches for predictive translation between animal models and humans.^20–23^ Translatable Components Regression (TransComp-R) was originally developed to integrate disparate multi-omics data from mouse models and humans to predict drug resistance phenotypes.^24^ What distinguishes TransComp-R from descriptive cross-species comparisons is the explicit goal of finding biological features and signals in the animal model that are predictive of phenotypes or outcomes in humans.

TransComp-R has been successfully applied to cross-species prediction of neurodegenerative disease biology^25–27^ and for assessment of novel therapeutic hypotheses in colitis.^28, 29^ In this study, we evaluated the cross-species translatability of cartilage transcriptomics data in two common models of mouse post-traumatic OA, surgical destabilization of the medial meniscus (DMM) and noninvasive anterior cruciate ligament rupture (ACLR). We hypothesized that transcriptomic features in each mouse model would be predictive of different aspects of human OA and leveraged TransComp-R to assess translatability. Furthermore, we extended TransComp-R to incorporate human variables of age and sex into mouse-to-human translation modeling to assess how these critical factors influenced the predictive capacity of our cross-species translation model.

## METHODS

### Study Selection and Gene Expression Data Processing

We searched the Gene Expression Omnibus data repository^30^ (accessed November 2024) for “osteoarthriti*”, restricting our search to *Homo sapiens* and *Mus musculus* records and using “*” as a wildcard character. From the search results, we then manually curated studies to those with mRNA expression for n ≥ 10 per group from human OA patients and n ≥ 3 per time point and experimental group within murine OA models. We further narrowed the datasets to those with controls and excluded datasets from genetic knockout mouse models of OA (**Figure 1**).

**Figure 1.**
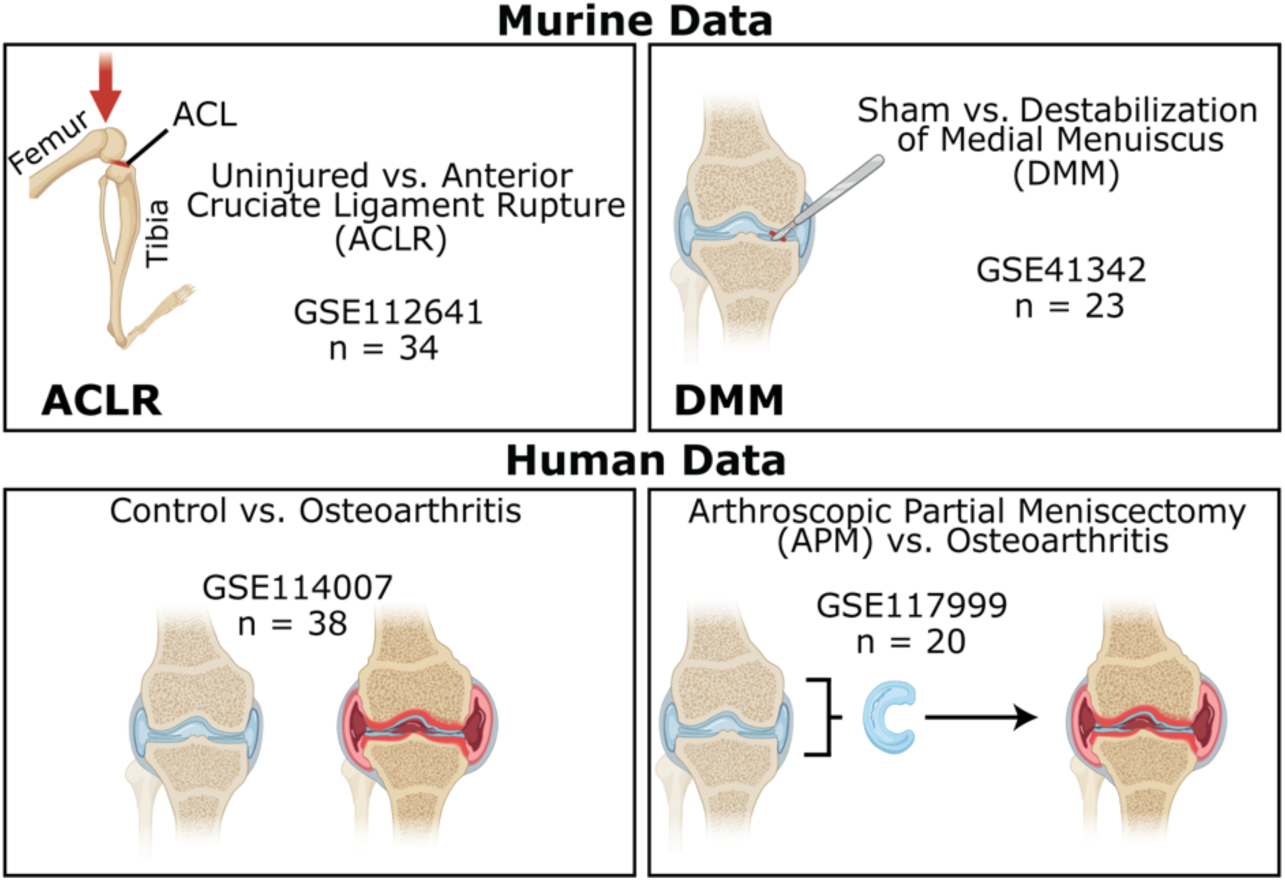
Selected murine and human datasets. Four separate datasets were selected from GEO, a publicly available data repository. The murine data contains transcriptomic data of control and ACLR models (GSE112641) and sham with DMM models (GSE41342). The human transcriptomic data contains subjects with Control and OA conditions (GSE114007) as well as APM compared to OA (GSE117999). Demographic variables of Sex and Age are included in the human dataset. The schematics of anatomy are from BioRender.com.

We selected human OA datasets (**Supplemental Table S1**) from two studies of cartilage retrieved during total knee replacement (GSE117999,^31^ n = 20; GSE114007,^32^ n = 38). Human dataset GSE114007 (hereafter “H007”) included gene expression for control donated cartilage and OA cartilage, and GSE117999 (hereafter “H999”) included gene expression for cartilage from arthroscopic partial meniscectomy (APM) and OA patients. Association of APM with increased risk of OA^33^ prevents H999 APM controls from being classified as healthy, motivating separate analysis of the two human datasets.

From the mouse datasets (**Supplemental Table S2**), we selected one DMM study (GSE41342,^34^ n = 23) and one ACLR study (GSE112641,^35^ n = 34). Murine genes were converted to human gene IDs with the Mouse Genome Informatics database, and both human and mouse datasets were collapsed to include only the overlapping one-to-one homolog genes for analysis.^36^ Baseline differential gene expression similarity was assessed using the Bioconductor tools *limma* (v3.58.1) and *edgeR* (v4.0.16) in R,^37^ controlling for time in mouse data and both Sex and Age in human data (control vs. disease, adjusted p < 0.05).

### Translatable Components Regression

Cross-species TransComp-R^24^ involves (1) construction of a principal component analysis (PCA) model on animal data, (2) creation of a cross-species translational space through projection of human data onto the principal component (PC) or latent variable representation of animal model data, and (3) construction of a regression model to relate the projected human data positions in translational space to human phenotypic variables (**Figure 2A**). This process helps identify the PCs of the animal data that best explain the phenotype from the human observations.

**Figure 2.**
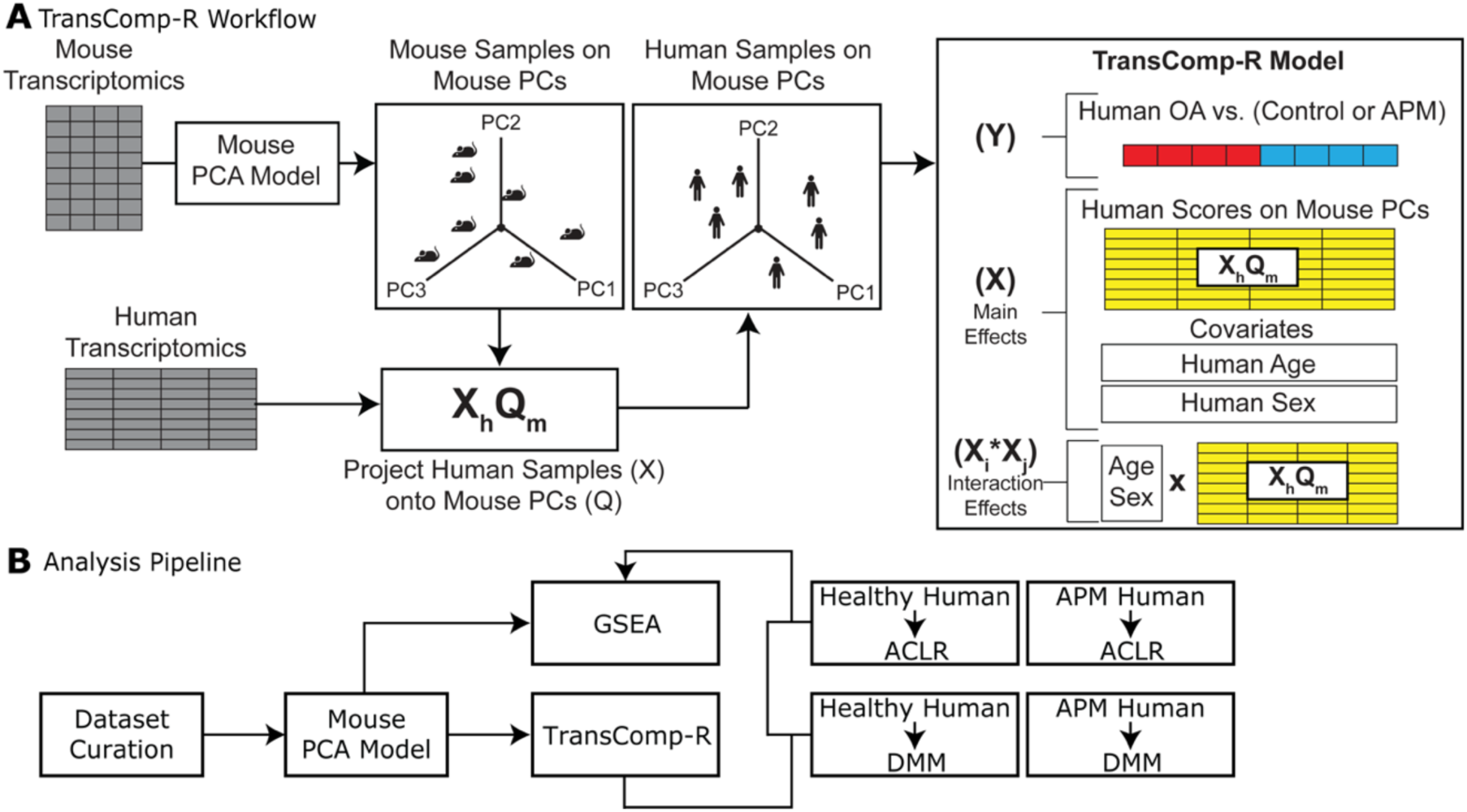
Analysis pipeline from principal components analysis (PCA) to translatable components regression (TransComp-R) model. (A) Mouse and human transcriptomics data are first matched for homologs. Human data is then projected into a constructed mouse PCA space. The synthesized human-mouse data containing genes and PCs are regressed against human OA condition using PC main effects and interaction effects of Age and Sex. PCs that significantly stratify OA and control status are selected for downstream analysis. (B) After data curation, the TransComp-R model is performed using ACLR and DMM pairs of healthy and APM human data. The loadings from the selected DMM and ACLR mouse PCs are analyzed through GSEA for biological interpretation.

We analyzed four TransComp-R models, mapping the DMM or ACLR datasets against human datasets H999 or H007 (**Figure 2B**), denoting these as DMM-H999, DMM-H007, ACLR-H999, and ACLR-H007. For each mouse-human comparison, the source PCA model was initially constructed using the DMM or ACLR dataset, each of which included controls, discarding PCs that explained less than 1% total variance. Next, Z-score normalized human data from either H999 or H007 was projected into the mouse PC space through matrix multiplication by mouse PC loadings. Human projections in mouse space were regressed against human OA status using *fitglm* (MATLAB, v9.13, The Mathworks, Natick, MA) to identify mouse PCs predictive of human disease. The significant mouse PCs identified by the regression modeling were analyzed through gene set enrichment analysis (GSEA).

### Incorporation of Human Clinical Covariates in Cross-species Modeling

We extended TransComp-R to account for human covariates of Age and Sex at the regression model stage to account for how these variables influence the predictive translatability of the mouse data. The incorporation of human clinical factors, and their interaction effects increased model complexity and necessitated down-sampling of the mouse PCs for analysis. We down-sampled the mouse PCs by building principal component regression models of the mouse data, relating mouse PCs to mouse OA status and time points, to identify predictive mouse PCs with regression coefficients providing p < 0.05. Using these significant PCs, we constructed TransComp-R models by setting up generalized linear models with increasing levels of complexity to assess how the predictive power of mouse PCs changed when human covariates and their interactions with mouse PCs were incorporated (**Table 1**).

**Table 1.** TransComp-R generalized linear model equations. PCX and PCY are placeholders for predictive PCs determined from the mouse PC regression as predictive of OA status in mouse data. The disease state is defined in each model as the OA status in the human dataset. Human covariates of Age and Sex are considered in all models but the “PCs Only” model, and interactions (×) among PCs and covariates are modeled in the Age, Sex, and Full models.

| Model | Equation |
| --- | --- |
| PCs Only | $Disease \sim PCX + PCY$ |
| Main | $Disease \sim PCX + PCY + Age + Sex$ |
| Age | $Disease \sim PCX + PCY + Age + Sex + (PCX \times Age) + (PCY \times Age)$ |
| Sex | $Disease \sim PCX + PCY + Age + Sex + (PCX \times Sex) + (PCY \times Sex)$ |
| Full | $Disease \sim PCX + PCY + Age + Sex + (PCX \times Age) + (PCY \times Age) + (PCX \times Sex) + (PCY \times Sex)$ |

### Pathway Enrichment Analysis of Significant PC Loadings

Pre-ranked PC loadings from ACL and DMM datasets were analyzed with GSEA in R (fgsea v1.28.0 and clusterProfiler v2.1.6).^38, 39^ KEGG (C2) and Hallmark (H) from the Molecular Signatures Database (msigdbr v7.5.1)^40^ were used to identify biological pathways.^41, 42^ The parameters for GSEA were established at 5 minimum and 500 maximum gene sets, and the tuning constant, epsilon, was set to 0. The default of 1000 permutations were used for the model. A Benjamini-Hochberg adjusted p-value less than 0.25 was used to identify potentially relevant pathways, whereas an adjusted p-value less than 0.10 was defined as significant.

### Cross-species Variance Explained Analysis

Because the variance a PC explains in mouse data may differ from the variance it explains in human data, we calculated the variance explained by a mouse PC *q_i_* in a human dataset (**Equation 1**) as:

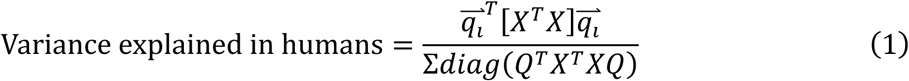

where *Q* are the mouse PC loadings, *X* indicates Z-score normalized human data, and *^T^*denotes the transpose of a matrix.

### Data and Core Availability

All data necessary to reproduce these analyses are publicly available through Gene Expression Omnibus with accession numbers GSE117999, GSE114007, GSE41342, and GSE112641. All code used for the analyses is publicly available at https://github.com/Brubaker-Lab/TOAST.

## RESULTS

### Baseline Signatures and Similarities of Mouse and Human OA Cartilage Transcriptomics

We first assessed the baseline similarity of our selected mouse DMM or ACLR datasets and H007 human OA vs. Control dataset or H999 OA vs. APM dataset via differential expression analysis (**Figure 3A**). 2,644 genes were differentially expressed within any dataset, with the DMM model having more differentially expressed genes (DEGs) in common with human OA than the ACLR model. Only one DEG, *IKBIP* (inhibitor of nuclear factor kappa-B-kinase-interacting protein), was common to all mouse models and human OA comparisons. In the human data, only 221 DEGs distinguished APM from OA in H999, but 950 DEGs were found between OA and Control in H007. DMM shared 24 DEGs with H999 and 143 DEGs with H007. In contrast, ACLR shared only 6 DEGs with H999 and 74 DEGs with H007. Based on these standard differential expression analysis metrics, the OA-associated gene signatures in DMM and ACLR models were more similar to the H007 OA vs. Control contrast. DMM captured more of the gene expression changes in the H999 APM vs. OA contrast than ACLR.

**Figure 3.**
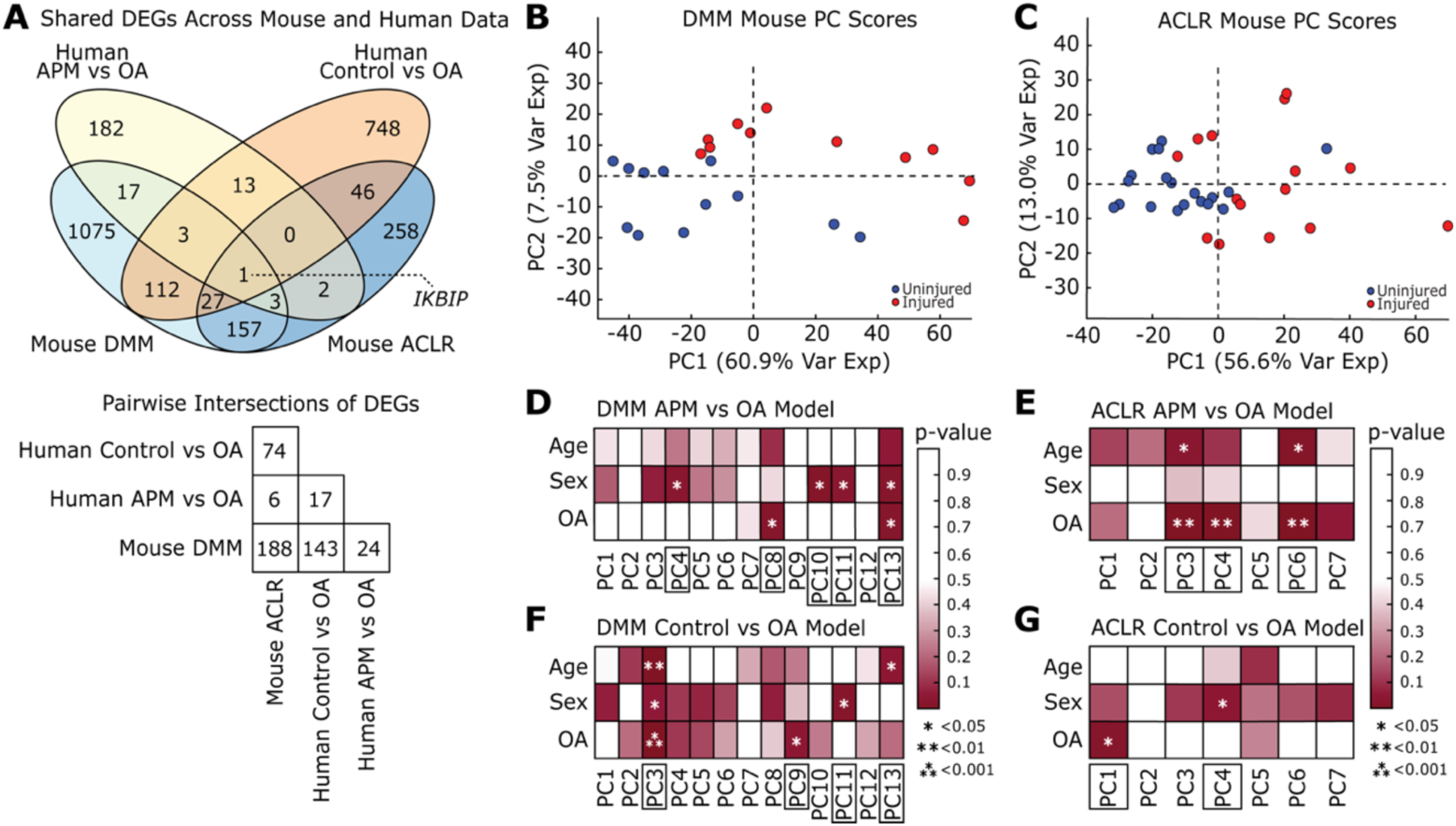
Baseline Signatures of Mouse and Human OA Cartilage Transcriptomes to Translatable Latent Variables from Mouse and Human Data. **(A)** Shared DEGs across human APM vs OA, human control vs OA, mouse DMM and mouse ACLR. Intersections of shared genes across the four datasets. **(B)** Principal component plots of PC1 and PC2 with the DMM mouse group **(C)** Principal component plots of PC1 and PC2 with the ACLR mouse group. **(D)** DMM Control vs OA models (DMM-H007) on Age, Sex, and OA conditions. **(E)** ACLR Control vs OA models (ACLR-H007) on Age, Sex, and OA conditions. **(F)** DMM APM vs OA models (DMM-H999) on Age, Sex, and OA conditions. **(G)** ACLR APM vs OA models (ACLR-H999) on Age, Sex, and OA conditions.

### Down-Sampling Translatable Latent Variables from Mouse and Human Data

We constructed PCA models for each mouse dataset and excluded PCs that explained less than 1% total variance. The DMM PCA model included 286 genes (FDR q < 0.05, 13 PCs, 93.9% variance explained) associated with DMM surgery status (**Figure 3B**). The ACLR PCA model included 486 genes (FDR q < 0.05, 7 PCs, 88.3% variability explained) associated with ACLR injury status (**Figure 3C**). Both mouse PCA models stratified the OA vs. control conditions in the mouse data alone, but a greater proportion of total variance is encoded on fewer PCs in ACLR data compared to DMM.

Next, we constructed principal component regression models to identify the mouse PCs predictive of joint injury status and any time-dependent behavior in the mouse PCs. For the DMM model, PC1, PC2, and PC7 were associated with both mouse time points and injury status (**Figure S1A**), For the ACLR model, PC1, PC3, and PC7 were associated with both time and injury status, and PC4 associated with injury status alone (**Figure S1B**).

After projecting Z-score normalized human gene expression data from H999 and H007 into each of the mouse PC spaces, we constructed TransComp-R regression models relating human projections in each space to human OA status, Age, and Sex. In total, twelve preliminary TransComp-R models were tested (**Figure 3D-3G**), including three human dependent variables (Age, Sex, and OA Status) and four cross-species pairs (ACLR-H007, ACLR-H999, DMM-H007, and DMM-H999).

When using the DMM mouse PCs to predict H007, we found PC3 was associated with human Age, Sex, and OA status while PC9, PC11, and PC13 were individually associated with human OA status, Sex, and Age, respectively (**Figure 3D**). For DMM-H999, only DMM mouse PC8 and PC13 were associated with human OA vs. APM status, while PC4, PC10, PC11, and PC13 were associated with human Sex (**Figure 3F**). Although DMM PC11 was also associated with Sex, PC11 explained less variance than PC10 and was only significant in the Sex model. We therefore only selected DMM PC4, PC8, PC10, and PC13 for the further DMM-H999 analysis. Despite starting with the same set of genes and parent PCA model, only the low-rank PC13 was consistently predictive of OA×Age status across H999 and H007. Critically, none of the PCs implicated as predictive of DMM mouse injury status by the mouse data-only PC regression model was later identified as predictive in the cross-species models, indicating that human signature translatability is not one-to-one for DMM.

For ACLR predicting H007, we identified PC1 as predictive of OA vs. Control status and PC4 as predictive of human Sex (**Figure 3E**). For ACLR-H999, PC3 and PC6 were predictive of human OA vs. APM status and Age, with PC4 being predictive of just OA vs. APM status (**Figure 3G**). No ACLR PCs predicted both OA vs. Control and OA vs. APM status. Unlike DMM, there were mouse PCs that were predictive of both ACLR injury status in mice and human OA vs. Control (PC1) and OA vs. APM (PC3, PC4), indicating that the ACLR mouse transcriptomics contained one-to-one translatable signatures of human OA status.

### Functional Analysis of Candidate Translatable Mouse Principal Components

We analyzed the pre-ranked loadings on the predictive PCs using GSEA to identify Hallmark pathways associated with the translatable DMM (**Figure 4A**) and ACLR (**Figure 4B**) mouse PCs. The identified DMM and ACLR mouse PCs encoded distinct axes of human-translatable OA biology. The ACLR model contained 884 significant genes whereas the DMM model had 2,100 significant genes. Among the Hallmark pathways enriched in DMM, only three pathways were significantly enriched (Benjamini-Hochberg p adj < 0.10), including Protein Secretion on DMM PC3 (p adj = 0.0635), E2F Targets on PC4 (p adj = 0.0004), and IL2 STAT5 signaling on PC8 (p adj = 0.0208). All other pathways were associated with cellular signaling, immune response, and metabolic pathways. Among ACLR PC1, we identified Apical Junction (p adj = 0.0370) associated with OA. On ACLR PC4, Allograft Rejection (p adj = 0.0004) and TNFα Signaling via NF-κB (p adj = 0.0860) were associated with non-OA conditions. Shared across the ACLR PCs, Epithelial Mesenchymal Transition was consistently and significantly associated with OA in three of the four selected ACLR PCs:PC1 (p adj = 0.0422), PC3 (p adj = 0.0034), and PC4 (p adj = 0.0641). Inflammatory Response was found to be significantly enriched on ACLR PC3 (p adj = 0.0838) and PC4 (p adj = 0.0049), with opposite normalized enrichment score directionality.

**Figure 4.**
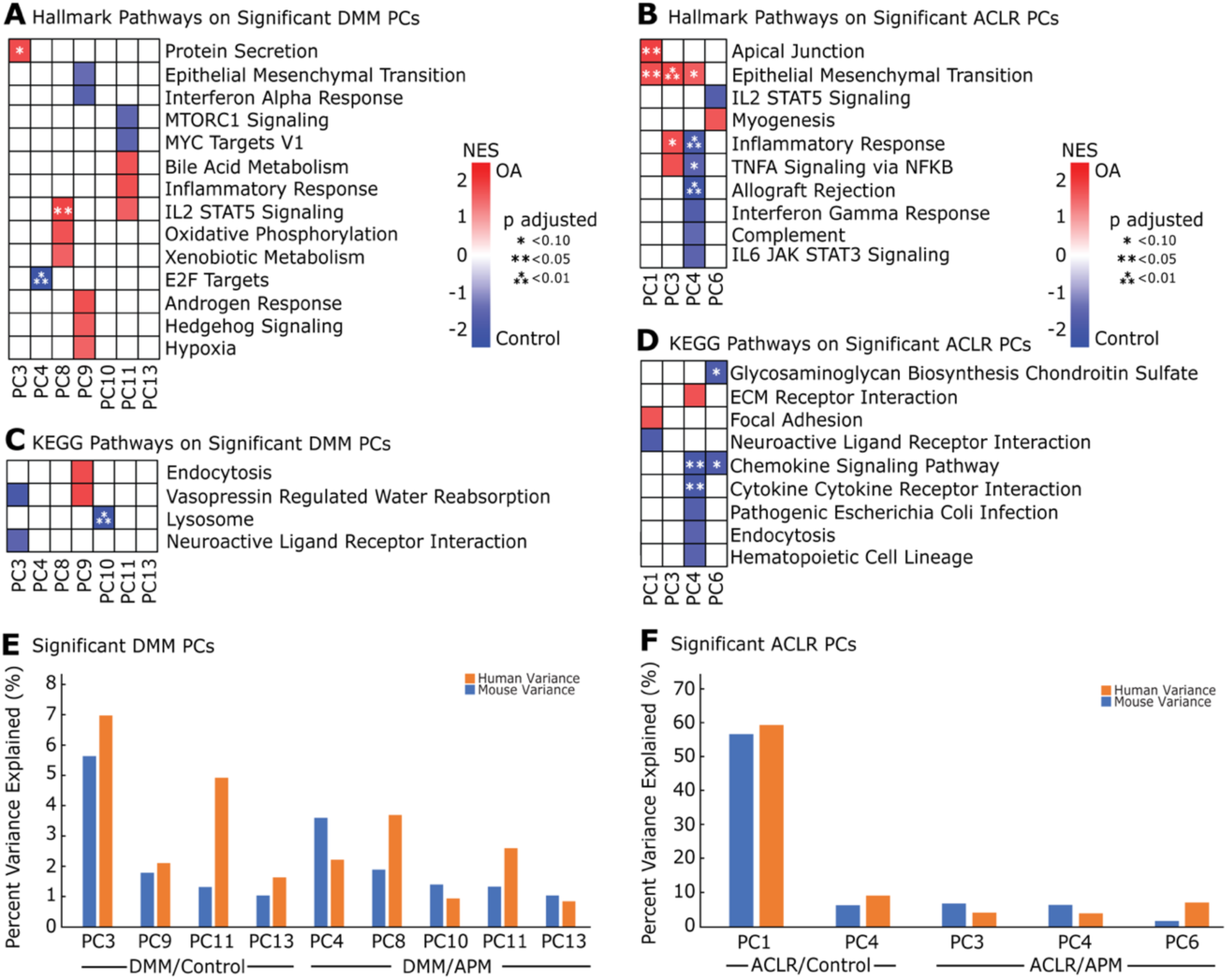
Pathways and Molecular Signatures Associated with Significant PCs from ACLR and DMM Datasets. **(A)** Hallmark pathway on significant PCs on the DMM group. **(B)** ACLR group. **(C)** KEGG pathways on DMM and **(D)** ACLR conditions. Normalized Enrichment score directions were referenced such that positive values were associated with OA whereas negative values were associated with non-OA conditions. Pathways denoted with * indicate significance (adjusted p < 0.10). No * indicates an adjusted p-value less than 0.25. **(E)** Percent variance explained significant DMM PCs in both human and mouse. **(F)** Percent variance explained of significant ACLR PCs in both human and mouse.

We also ran pre-ranked loadings using the KEGG pathway for both DMM (**Figure 4C**) and ACLR PCs (**Figure 4D**). Of the pathways enriched in DMM, only four had an adjusted p-value less than 0.25: Endocytosis, Vasopressin Regulated Water Reabsorption, Lysosome, and Neuroactive Ligand Receptor Interaction. Of these pathways, only Lysosome was significant (p adj = 0.0079), with associations to non-OA conditions. From the ACLR PCs, general pathways included immune signaling, where the Chemokine Signaling Pathway was significant on ACLR PC4 (p adj = 0.0253) and PC5 (p adj = 0.0278), with Cytokine-Cytokine Receptor Interaction also significant on PC4 (p adj = 0.0631). Additionally, on ACLR PC4, Glycosaminoglycan Biosynthesis Chondroitin Sulfate was significantly enriched for non-OA conditions (p adj = 0.0631).

In both DMM and ACLR, the human variance explained was often higher than the mouse variance explained. Some translatable PCs explained greater variability in human OA than in mouse DMM (**Figure 4E**). Significant DMM PCs explained no more than 8% variance in human and mouse (**Figure 4F**). Explained human variance was also higher than for mouse among the ACLR PCs. PC1 explained more than 50% variance, whereas the remaining significant PCs explained less than 10% variance. The result that mouse PCs implicated as translatable to humans explain a higher proportion of human data variability compared to their explanatory power in mouse data indicates that latent, hidden signals in mouse models may be predictive of human biology despite low explanatory power for mouse biology.

### Covariate-Conditioned Cross-Species Modeling of DMM Mouse Cartilage Transcriptomics

Iteratively building up model complexity and incorporating different covariates and interactions enabled the identification of consistent associations of mouse PCs with human OA disease status while controlling for patient characteristics (**Table 1**). With DMM comparison to H999, we identified PC8, PC13, and human Age as being significantly predictive of human OA vs. APM status when controlling for Age and Sex, along with a significant interaction between PC8 and Age (p = 0.047) (**Figure 5A**). Plotting the projections of the human samples on the predictive mouse PCs (**Figure 5B**) and PC8 vs. Age (**Figure 5C**) revealed a separation of OA samples vs. APM that tracked with Age.

**Figure 5.**
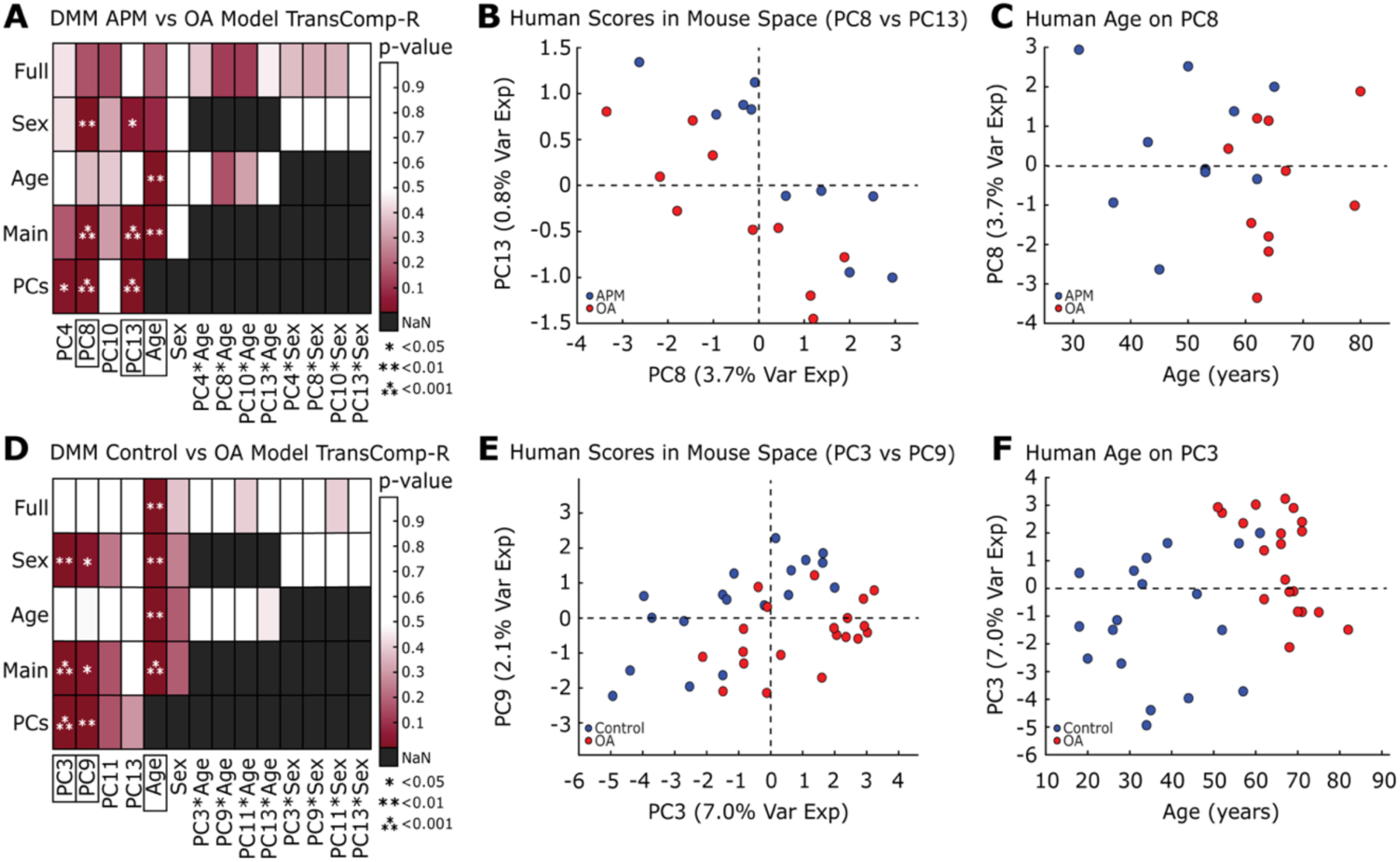
Generalized Linear Equation Modeling of DMM PCs from TransComp-R. **(A)** DMM PC8, PC13, and Age are selected from the DMM APM vs OA model. **(B)** Principal component plot of DMM PC8 and PC13. **(C)** DMM PC8 scores were visualized with respect to human age. **(D)** DMM PC3, PC9, and Age are selected from the DMM Control vs OA Model. **(E)** DMM PC3 and PC9 scores plot. **(F)** DMM PC3 scores are separated by human age. All percent variance explained is in humans.

Modeling the DMM mouse with H007, we found PC3, PC9, and human Age were associated with the human OA vs. Control contrast (**Figure 5D**), with no statistical interactions between human clinical variables and mouse PCs. Plotting human scores on PC3 and PC9 (**Figure 5E**) and on PC3 vs. Age (**Figure 5F**) revealed stratification by OA status and tracking of Age with OA status.

### Covariate-Conditioned Cross-Species Modeling of ACLR Mouse Cartilage Transcriptomics

The ACLR-H999 TransComp-R model identified PC4 and Age as predictive factors for the OA vs. APM contrast (**Figure 6A**). An age-dependent separation of human OA vs. APM was observed with PC4 (**Figure 6B**). The ACLR-H007 TransComp-R model revealed patterns of PC1, Age, and a statistical interaction between ACLR PC1 and human Sex as significantly predictive of OA vs. Control status (**Figure 6C**). PC1 separated OA and Control in an age-dependent manner (**Figure 6D**). Interestingly, we identified a Sex-dependent separation of human OA vs. Control along ACLR mouse PC1, where the magnitude of separation along PC1 was significantly greater for female compared to male humans (**Figure 6E**), showing that TransComp-R was able to detect human sex-specific, translatable signatures. This identification of ACLR mouse PC1 as predictive of human OA status, along with PC1’s predictive power for mouse injury status, indicates that the gene signatures, pathways, and biological variability encoded on this mouse PC represents a potential direct mapping between the ACLR mouse biology and human OA.

**Figure 6.**
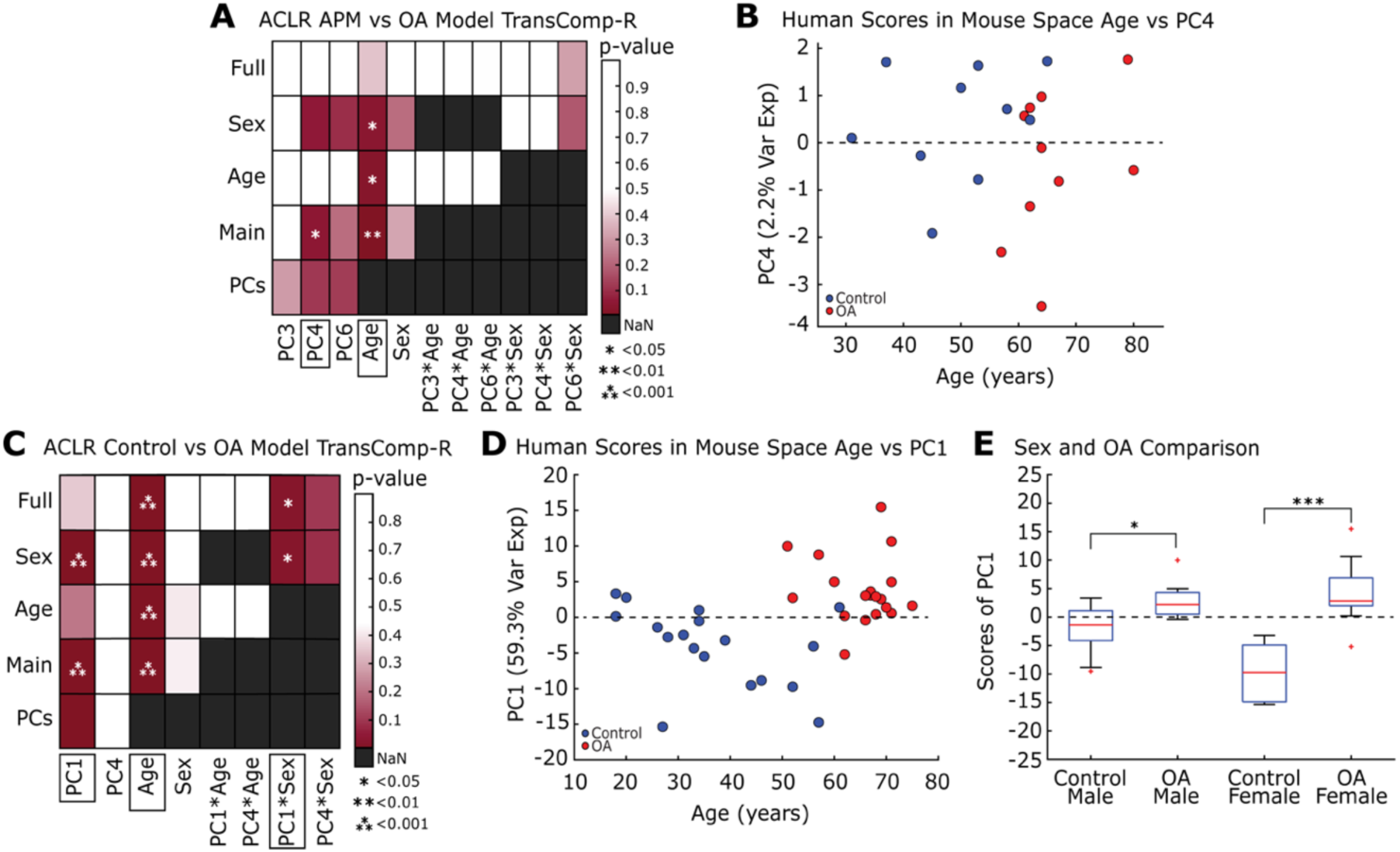
Generalized Linear Equation Modeling of ACLR PCs from TransComp-R. **(A)** ACLR PC4 and Age were selected from models of ACLR APM vs OA. **(B)** Scores plot of PC4 against human age. **(C)** ACLR PC1, Age, and PC1*Sex were significant in the ACLR Control vs OA models. **(D)** Scores plot of PC1 against human age. **(E)** Comparison of PC1 scores separated by sex and OA condition. All percent variance explained is in humans.

## DISCUSSION

In this study, we evaluated the translatability of two commonly used mouse OA models into different cartilage transcriptomic features of human OA using TransComp-R, a framework that enables cross-species prediction of pathobiology. Pathway analysis enabled the identification of mouse transcriptional signatures translatable to human OA. At over 50% variance explained for human OA status along PC1, the ACLR mouse model was more translatable to humans. Inclusion of human covariates of Age and Sex altered the identified translatable features from both mouse models, enabling the identification of a candidate sex-specific effect predictive of human OA status for the ACLR mouse model. Covariate-aware TransComp-R provides a path to incorporate human sex as a biological variable in assessing sex-based variation from mouse studies and can model a wide range of human covariates in assessing the translatability of mouse studies.

### Mouse to Human Conversion and Differential Expression

Significant PCs representing variance in human data projected onto murine principal component loadings were identified through generalized linear modeling. Both DMM-H007 and DMM-H999 models contained four significant PCs, with a cumulative variance explained in human data of 15.61% (PCs 3, 9, 11, 13) and 7.64% (PCs 4, 8, 10, 13), respectively. ACLR models had fewer significant components (two significant PCs with ACLR-H007; three significant PCs with ACLR-H999), but the cumulative variance explained (68.47% in ACLR-H007 PCs 1, 4; 15.09% in ACLR-H999 PCs 3, 4, 6), was greater than the variance explained in DMM models. This distinction of ACLR models for fewer PCs explaining greater variance is also stronger in the prediction of human control vs. OA status than APM vs. OA status, suggesting better translatability where there is similarity in the experimental control groups. The H007 dataset compared transcriptomic profiles from healthy cartilage obtained through a tissue bank to those of OA cartilage.^32^ Similarly, for noninvasive ACLR, the sham joint is not injured nor opened ^8^. Although ACLR data was found to be more directly translatable to human OA data, DMM and potentially other surgical models are likely still applicable to human disease under certain circumstances. Association of human APM vs. OA to murine surgical sham vs. DMM presented an interesting comparison, since both non-OA groups are associated with a prior history joint disruption. A comparison of the synovium of sham and DMM mice found almost 400 dysregulated miRNAs at the 1- and 6-week timepoints in both experimental groups,^43^ suggesting that arthrotomy alone accounts for some changes detected in surgical models of OA. Synovial inflammation and changes to cartilage and bone morphology have been previously identified in sham-operated joints.^44, 45^ Similarly, the H999 dataset used cartilage from patients requiring arthroscopic resection of meniscal injury as a comparison group.^46^ Because a meniscal tear already disrupts the joint, molecular signatures associated to the injury may still be present in these patients. Patients requiring APM generally have persisting symptoms that require surgical management,^47^ suggesting progressive meniscus injury and potentially early OA.^48^ Indeed, fewer DEGs (221) were found between APM and OA in H999 than between OA and control in H007 (950), suggesting that injury-associated gene signatures in APM may overlap with those associated with OA. The human APM group may thus be more akin to surgical sham in the DMM studies or even early mouse OA time points. Therefore, beyond representing the variability of human disease, the two human datasets selected for this study fortuitously parallelled some aspects of the experimental design of DMM and ACLR models.

### Modeling Human Covariates using TransComp-R on ACLR and DMM

Both the DMM^34^ and ACLR^35^ datasets show potentially translatable biology when evaluated using TransComp-R. All four TransComp-R comparison models identified human Age as a significant covariate in the prediction of human disease, supporting that the model was consistent with a prominent risk factor for OA.

The ACLR-H007 TransComp-R model indicated that the interaction of ACLR PC1 and human sex was significantly predictive of human OA status. This finding was striking because the mouse ACLR data used to train the underlying PCA model did not include mouse sex, underscoring the importance of reporting sex with OA murine studies. Further comparison, by Mann-Whitney test, showed that ACLR PC1 values were significantly different between human males and females in both control and OA groups. Covariate-aware TransComp-R provides a path to incorporate human Sex as a biological variable in assessing mouse studies, whether or not mouse sex is reported, and can model a wide range of other human covariates in assessing the translatability of mouse studies.

### Gene Set Enrichment Analysis of Translatable Murine Signatures

The pathways enriched in ACLR conditions differed those identified in the DMM model. We identified biological processes related to cell signaling and metabolism from the DMM mouse PCs, including Protein Secretion, IL2 STAT5 Signaling, and Lysosome. The association to protein secretion is supported by a study that found differentially expressed proteins from human OA cartilage secretome, including growth-regulated alpha protein, stromelysin-1, LYR motif-containing protein 5, and Ig alpha-1 chain C region.^49^ A separate computational study also found IL2 STAT5 Signaling to be enriched from GSEA when comparing OA and healthy synovial tissue.^50^ Lysosomal dysregulation was found to contribute to chondrocyte apoptosis in both OA and aged cartilage through mitochondrial damage.^51^

ACLR mouse PCs were strongly associated with immune function and cellular pathways. We identified Epithelial Mesenchymal Transition as an enriched pathway in OA, and others have found this pathway is associated with synovial inflammation and cartilage destruction.^52^ Immune-associated pathways such as Chemokine Signaling and Inflammatory Response were also involved in conditions of OA.^53, 54^ A computational study also found sets of genes enriched for Allograft Rejection associated with OA.^50^ Lastly, Glycosaminoglycan Biosynthesis Chondroitin Sulfate serves an important role in joint health, and dysfunction of this pathway may precede OA development.^55^ Further investigation of these translatable pathways may provide insight in OA progression across DMM and ACLR models.

### Outlook and Conclusions

Expansion of our TransComp-R approach can integrate other experimental or clinical information that may enhance model accuracy and robustness. Incorporating OA-associated serum biomarkers,^56, 57^ race,^58, 59^ and other clinical covariates may better account for heterogeneity in human OA, improving overall understanding of the disease. Integration of other omics data could also uncover additional layers of cross-species translatable OA biology, including proteomics, ^60^ metabolomics,^61^ and even microbiome.^62^ However, open access availability of multi-omics experimental data in human OA and mouse models is a critical barrier to expanding our TransComp-R approach.

In our use of TransComp-R to analyze data from two established mouse OA models, we identified gene signatures that predicted human OA. Despite being limited to the currently available open access datasets and homologous gene pairs across mouse and human cartilage transcriptomic data, we found that ACLR better predicted human OA status, particular in the contrast between donor control cartilage and OA tissues obtained prior to joint replacement surgery. Because murine models with different methods of OA induction differentially translated to human OA status, caution should be exercised when selecting an animal model to represent human disease progression or assess treatments. Ensuring that preclinical models best reflect human outcomes could improve our understanding of translatable pathologies and the rate of success in translating such findings to human therapeutics and interventions. Working towards these goals, we demonstrated here that different mouse OA models, DMM and ACLR, encoded distinct axes of translatable biology that relied on human age as a risk factor and sex as a biological variable.

## AUTHOR CONTRIBUTIONS

**Conception and Design:** MF, MP, DKB, and DDC. **Data Acquisition:** MF, BKB, MP, and SRD. **Data Analysis and Interpretation:** MF, BKB, MP, DKB, and DDC. **Writing Original Article:** MF, BKB, MP, DKB, and DDC. **Writing-Review and Editing:** MF, BKB, DKB, and DDC. **Funding Acquisition:** DDC. **Project Administration:** DDC. **Final Approval of Article:** MF, BKB, MP, SRD, DKB, and DDC.

