## Supplementary material for "Computational Translation of Mouse Models of Osteoarthritis Predicts Human Disease": Table S1

**Supplemental Table S1. Human data sets with relevant parameters considered for study inclusion.** The Gene Expression Omnibus was searched for gene expression data from joint tissues at the late to end stages of osteoarthritis (OA) with $n \geq 10$. All OA tissues were obtained at the time of total knee arthroplasty (TKA), which can be considered end-stage OA. Control tissue sources included tissue banks and arthroscopic partial meniscectomy (APM). Two datasets (GSE114007, GSE117999) were chosen for analysis because non-OA controls were available for cartilage in these data. The data sets selected are bolded in the table below.

| Dataset | Tissues and Source | n | Age Range | Sex | Platform(s) |
|---|---|---|---|---|---|
| GSE98460 | Medial, Lateral Tibial Plateau Cartilage, TKA | 46 | 51-72 years old (mean = 62.6) | | GPL16686 [HuGene-2_0-st] Affymetrix Human Gene 2.0 ST Array [1] |
| **GSE114007** | **Medial, Lateral Tibial Plateau Cartilage, TKA** | **20** | **52-82 years old (mean = 66)** | **12 F 8 M** | **GPL11154 Illumina HiSeq 2000, GPL18573, Illumina NextSeq 500 [2]** |
| | **Tissue Bank** | **18** | **18-61 years old (mean = 38)** | **5 F 13 M** | |
| **GSE117999** | **Cartilage, TKA** | **12** | **53-80 years old (mean = 65.3)** | **9 F 3 M** | **GPL20844 Agilent-072363 SurePrint G3 Human GE v3 8x60K Microarray 039494 (no citation was listed; however, this dataset appears in [3])** |
| | **Cartilage, APM** | **12** | **31-65 years old (mean = 49.2)** | **5 F 7 M** | |
| GSE98918 | Posterior horn of the medial meniscus, TKA | 12 | 53-80 years old (mean = 65.3) | 9 F 3 M | GPL20844 Agilent-072363 SurePrint G3 Human GE v3 8x60K Microarray 039494 [3] |
| | Posterior horn of the medial meniscus, APM | 12 | 31-65 years old (mean = 49.2) | 5 F 7 M | |

*NOTE: This bibliography refers only to literature cited in this table. Numbering does not correspond to the literature cited in the primary manuscript.*


1. Rai MF, Sandell LJ, Barrack TN, Cai L, Tycksen ED, Tang SY, et al. A Microarray Study of Articular Cartilage in Relation to Obesity and Severity of Knee Osteoarthritis. Cartilage 2020; 11: 458-472.
2. Fisch KM, Gamini R, Alvarez-Garcia O, Akagi R, Saito M, Muramatsu Y, et al. Identification of transcription factors responsible for dysregulated networks in human osteoarthritis cartilage by global gene expression analysis. Osteoarthritis Cartilage 2018; 26: 1531-1538.
3. Brophy RH, Zhang B, Cai L, Wright RW, Sandell LJ, Rai MF. Transcriptome comparison of meniscus from patients with and without osteoarthritis. Osteoarthritis Cartilage 2018; 26: 422-432.
