## Supplementary material for "Computational Translation of Mouse Models of Osteoarthritis Predicts Human Disease": Table S2

**Supplemental Table S2. Murine data sets with relevant parameters examined for study inclusion.** The Gene Expression Omnibus was searched for gene expression data from joint tissues from osteoarthritis (OA) models in mice, including destabilization of the medial meniscus (DMM), rupture of the anterior cruciate ligament (ACLR) and natural onset. Controls, including sham or contralateral joints, and genetic variations are indicated separately from wild-type mice in the OA groups. The GSE41342 dataset was chosen for DMM with surgical sham control, and the GSE112641 dataset was chosen for ACLR with contralateral control.

| Dataset | Model | Background & Genetics | Tissues Analyzed | Age at Injury | Time Points (n, ea. timepoint) | Sex | Data Type |
|---|---|---|---|---|---|---|---|
| GSE93008 | DMM | C57BL6 | Medial tibial cartilage and subchondral bone | 10-12 weeks | 1-, 6- weeks post (n=4) | M | miRNA [1] |
| | DMM | Jaffa | | | 1-, 6- weeks post (n=3) | | |
| | Sham | C57BL6 | | | 1-, 6- weeks post (n=4) | | |
| | Sham | Jaffa | | | 1-, 6- weeks post (n=3) | | |
| GSE53857 | DMM | C57BL6 | Medial tibial cartilage | 10 weeks | 2-, 4-, 8- weeks post (n=3) | M | Microarray [2] |
| | Contra. | C57BL6 | | | 2-, 4-, 8- weeks post (n=3) | | |
| GSE45793 | DMM | C57BL6 | Cartilage | 10 weeks | 1-, 2-, 6-weeks post (n=4 each) | M | Microarray [3] |
| | DMM | C57BL6 Adamts5 Δcat | | | 1-, 6-weeks post (n=4 each) | | |
| | Contra. | C57BL6 | | | 1-, 2-, 6-weeks post (n=4 each) | | |
| | Contra. | C57BL6 Adamts5 Δcat | | | 1-, 6-weeks post (n=4 each) | | |
| GSE41342 | DMM | C57BL6 | Whole joint | 12 weeks | 0-, 2-, 4-, 8-, 16- weeks post (n=3) | M | Microarray [4] |
| | Sham | C57BL6 | | | | | |
| GSE33754 | Natural onset | CBA | Cartilage | N/A | 8-10, 18-20, 40-42 weeks old at sacrifice (n=5, 5, 9) | M | Microarray [5] |
| | Natural onset | STR/ort | | | 8-10, 18-20, 40-42 weeks old at sacrifice (n=6, 7, 14) | | |
| GSE26475 | DMM | C57BL6 | Whole joint | 10-12 weeks | 6 hours, 3 days, 7 days post (n=3) | M | Microarray [6] |
| | Sham | C57BL6 | | | 6 hours, 3 days, 7 days post (n=3) | | |
| GSE112641 | ACLR | C57BL/6N | Whole joint | 10 weeks | 1-, 7-, 14- days post (n=5) | M/F | RNA-seq [7] |
| | Contra | C57BL/6N | | | 1-, 7-, 14- days post (n=5) | | |
| | ACLR | STR/ort | | | 1-, 7-, 14- days post (n=5) | | |
| | Contra. | STR/ort | | | 1-, 7-, 14- days post (n=5) | | |

| | | | | | | | |
|---|---|---|---|---|---|---|---|
| | ACLR | MRL/MpJ | | | 1-, 7-, 14- days post (n=5) | | |
| | Contra. | MRL/MpJ | | | 1-, 7-, 14- days post (n=5) | | |
| GSE140785 | DMM | C57BL6 | | | 4-, 8-, 16-weeks post (n=3) | | |
| | Sham | C57BL6 | L3-L5 DRG | 10 weeks | 4-, 8-, 16-weeks post (n=3) | M | Mircoarray [8] |
| | Naïve | C57BL6 | | | 4-, 8-, 16-weeks post (n=3) | | |

*NOTE: This bibliography refers only to literature cited in this table. Numbering does not correspond to the literature cited in the primary manuscript.*
