## Supplementary material for "Computational Translation of Mouse Models of Osteoarthritis Predicts Human Disease": Figure S1

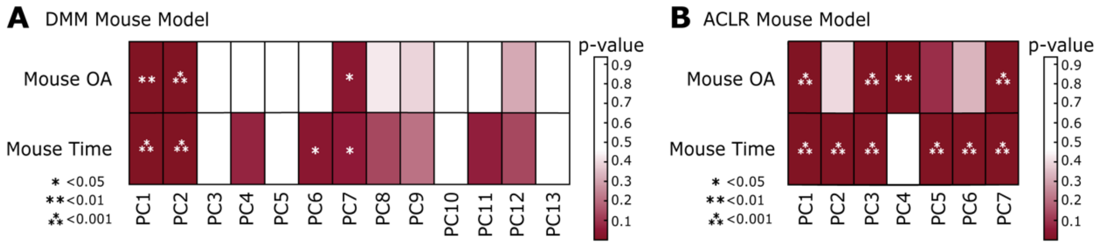


**Supplemental Figure S1. Selection of significant PCs from DMM and ACLR mouse models** **(A)** DMM mouse models on OA and Time status. **(B)** ACLR mouse models on OA and Time status.
